# DTSyn: a dual-transformer-based neural network to predict synergistic drug combinations

**DOI:** 10.1101/2022.03.29.486200

**Authors:** Jing Hu, Jie Gao, Xiaomin Fang, Zijing Liu, Fan Wang, Weili Huang, Hua wu, Guodong Zhao

## Abstract

Drug combination therapies are superior to monotherapy for cancer treatment in many ways when addressing tumor heterogeneity issue. For wet-lab experiment, screening out novel synergistic drug pairs is challenging due to the enormous searching space of possible drug pairs. Thus, computational methods have been developed to predict drug pairs with potential synergistic function. Notwithstanding the success of current models, the power of generalization to other datasets as wells as understanding of mechanism for chemical-chemical interaction or chemical-sample interaction are lack of study, hindering current algorithms from real application. In this paper, we proposed a deep neural model termed DTSyn (Dual Transformer model for drug pair Synergy prediction) based on multi-head attention mechanism to identify novel drug combinations. We designed a fine-granularity transformer for capturing chemical substructure-gene and gene-gene associations and a coarse-granularity transformer for extracting chemical-chemical and chemical-cell line interactions. DTSyn achieves highest Receiver operating characteristic area under curve (ROC AUC) of 0.73, 0.78. 0.82 and 0.81 on four different cross validation tasks, outperforming all competing methods. Further, DTSyn achieved best True Positive Rate (TPR) over five independent datasets. The ablation study showed that both transformer blocks contributed to the performance of DTSyn. In addition, DTSyn can extract interactions among chemicals and cell lines, which may represent the mechanisms of drug action. Thus, we envision our model a valuable tool to prioritize synergistic drug pairs by utilizing chemicals and transcriptome data.

## 1 Introduction

Drug combinations, compared to monotherapies, have the potential to increase efficacy, reduce host toxicity and adverse side effects and overcome drug resistance [Chou, 2006, O’Neil et al., 2016]. However, identifying novel synergistic drug combinations has been a laborious process and the vast number of possible drug pairs makes it difficult to screen them all by experiments. Though high-throughput screening (HTS) has been used to prioritize novel drug pairs, testing the whole complete combination space is still unfeasible [Goswami et al., 2015, Morris et al., 2016]. Thus, novel computational methodologies to facilitate the discovery of drug combination therapies are needed.

Recently, the release of large-scale datasets has enabled the exploring of machine learning based models or even neural networks on drug combinations. DrugCombDB has released 739,964 drug combinations [Zagidullin et al., 2019]. Also, transcriptome dataset of cancer cell lines, such as Cancer Cell Line Encyclopedia (CCLE) is available [Ghandi et al., 2019]. This project provides more than 1,000 cancer cell lines with comprehensive genetic characterizations across 39 cancer types. Indeed, many computational approaches for screening relevant drug combinations have been merged. For example, Preuer et al. proposed a deep learning model for predicting drug combination synergy scores by using both compound and genomic information as inputs [Preuer et al., 2018]. However, The omics data representing cell-line status was integrated with chemical inputs by simple concatenating operation, which lacks biological information and interpretability.

By taking biological interactions into considerations, Jiang et al. proposed a Graph Convolution Network (GCN) based model to prioritize potential synergistic drug pairs by performing heterogeneous graph message passing mechanism from a biological graph including drug and protein nodes [Jiang et al., 2020]. The computation method based on GCN was restricted to certain cell lines which limits the generalization of the model [Jiang et al., 2020]. Sun et al. presented a deep tensor factorization model which integrated tensor factorization method and canonical feed forward neural network, to predict drug synergy [Sun et al., 2020]. Furthermore, Menden et al. reported AstraZeneca’s drug combination dataset and results of a DREAM Challenge for predicting synergistic drug pairs [Menden et al., 2019]. These methods mentioned above only consider extracting chemical-cell line associations from one granularity, neglecting other dimensional interactions. To address the aforementioned problems, we proposed a dual-transformer based deep neural network named DTSyn (Dual-Transformer neural network predicting Synergistic pairs) for predicting potential drug synergies. As we all know, transformers [Vaswani et al., 2017] have been widely used in many computation areas including computer vision, natural language processing and even biological computing[Yuan et al., 2021, Parmar et al., 2018, Wolf et al., 2020, Jumper et al., 2021]. DeepCE utilized transformer, which can learn biological relations between chemical substructures and genes, to predict drug-induced expression profiles[Pham et al., 2021]. In this paper, we utilized two-branch transformer to capture biological associations between chemical-chemical, chemical substructure-gene, gene-gene and chemical-cell line. First, a graph convolution neural network (GCN) [Kipf and Welling, 2016] was applied to calculate the atom-level feature vectors of chemicals and it was designed to learn the substructure information of each drug. Second, the fine-granularity transformer block was used to capture relationships among chemical substructures, genes and gene-gene interactions. Specially, the gene feature vectors were obtained from a pre-trained model node2vec [Grover and Leskovec, 2016]. Meanwhile, the fine-granularity transformer block was designed to capture associations among both chemicals and cell line. Finally, multi-layer perceptron (MLP) was used to predict synergistic drug combinations from the updated features of chemicals and cell lines. DTSyn outperformed other comparative methods in four cross validation tasks and our model also showed best performance over five independent data sets. In addition, we explored the effectiveness of self-attention for extracting chemical substructure-gene interactions, gene-gene interactions and chemical-chemical interactions, and interestingly, we found that genes related to tumor proliferation, tumor metastasis, cell apoptosis and chemotherapy had much higher attention scores from the example of drug combination of ETOPOSIDE and MK-2206 in cell COAV3, which may help us to understand the mechanisms of drug combinations. In addition to known drug pair datasets, we validated our model using novel drug pair datasets. In summary, we believe that DTSyn could be an effective tool for identifying novel synergistic drug pairs.

## 2 Materials and methods

### 2.1 Synergy Data Collections

The Drug-Drug Synergy (DDS) data were obtained from O’Neil et al.’s work [O’Neil et al., 2016]. The DDS dataset contain 23,052 drug pairs, where each pair comprises two chemicals and a cancer cell line, covering 39 cancer lines across 7 different tissue types. The number of unique drugs was 38. There were 24 FDA-approved drugs and 14 experimental drugs [Preuer et al., 2018]. Replicating drug pairs were averaged as the final unique drug combinations. For noisy data removing and label balancing, we selected 10 as a threshold to classify the drug pair-cell line triplets. Triplets with synergistic score higher than 10 were positive, and those less than 0 were negative. Therefore, we obtained 13,243 unique triplets, consisting 38 drugs and 31 cell lines.

The independent test data includes AstraZeneca’s dataset [Menden et al., 2019], and the FLOBAK [Flobak et al., 2019], the ALMANAC [Holbeck et al., 2017], the FORCINA [Forcina et al., 2017] and the YOHE datasets [Zagidullin et al., 2019]. Four commonly-used criteria models were employed, including Loewe [Loewe, 1953], Bliss [Demidenko and Miller, 2019], HSA [Malyutina et al., 2019] and ZIP scores [Yadav et al., 2015]. Besides, Malyutina et al. utilized S score, which has been proved that it cloud measure the synergy level of drug combinations and predict the most synergistic and antagonistic drug pairs [Malyutina et al., 2019]. According to the above five criteria, we selected the pairs with all criteria greater than 0 as synergistic and those all less than 0 as antagonistic. Due to the limited expression profiles in corresponding cell lines, a total of 18813 combinations were obtained.

### 2.2 Expression profiles

The expression profiles of cancer cell lines were derived from Cancer Cell Line Encyclopedia (CCLE) [Ghandi et al., 2019]. The corresponding genes from LINCS L1000 project were extracted to represent the original cell line features [Subramanian et al., 2017].

#### 2.2.1 Framework of DTSyn

The framework of DTSyn presented in **Fig. 1**. The DTSyn model was constructed with two tracks: a fine-granularity block and a coarse-granularity block. There were four input modules including two chemical features, cell line expression profiles and gene embeddings. First of all, two chemicals were received by GCNs to extract the substructure information respectively. Integrating with pre-trained gene embeddings, the concatenated matrix were fed into the fine-granularity transformer encoder block. On the other hand, the expression profiles were encoded by MLP and concatenated with the pooled features following GCNs which generated inputs for the coarse-granularity transformer block. The embeddings from two transformer blocks were subsequently concatenated as the high-level feature propagating to the final prediction layer for classification of synergistic label.

**Figure 1:**
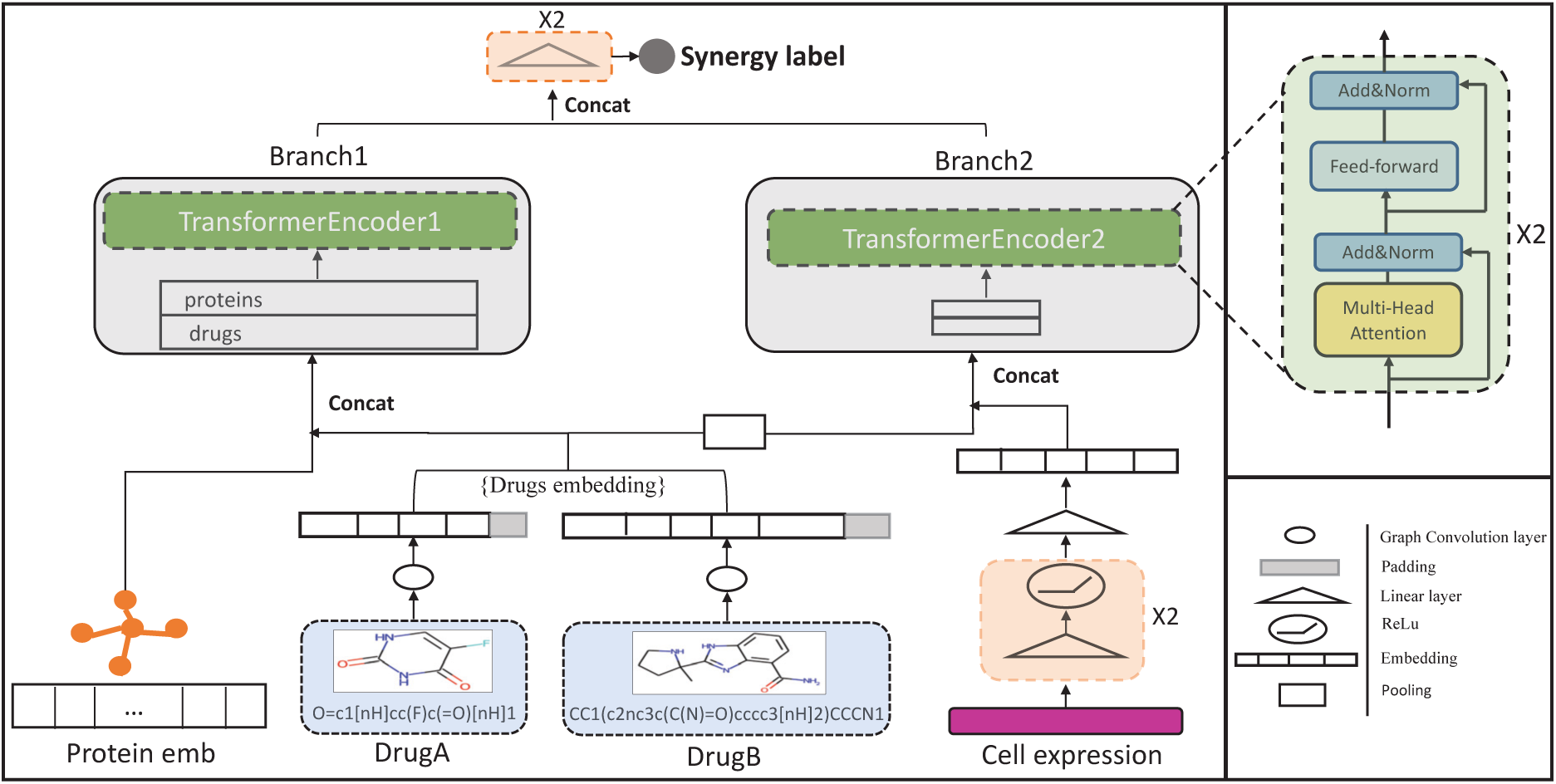
Overview of DTSyn. This model consists of two tracks which capture fine-granularity and coarse-granularity associations. The drugs features processed through GCN blocks concatenated with gene embeddings are fed into the fine-granularity transformer encoder block to learn the chemical substructures-gene interactions. Condensed cell line expression profiles processed by multi-layer preceptron (MLP) and pooled drugs features are used by the coarse granularity transformer encoder block, which can capture chemical-cell line and chemical-chemical associations. The final synergy label is obtained from the high-level features.

### 2.3 Drug features

In order to obtain the representation of chemical substructures, we used GCN model which trained a chemical graph structure as an input and updated vector embeddings of each atom from its neighbours [Pham et al., 2021]. Two-layer GCNs [Kipf and Welling, 2016] were chosen to make sure that each atom could see its two-hop neighbours. It indicated that the updated feature vector of each atom represented chemical substructures. We adopted RDKit [Landrum et al., 2013] to convert the SMILES [Weininger, 1988] format of chemicals to graphs. For the atom-level features of each chemical, we use DeepChem [Ramsundar et al., 2019] as initial atomic features, which were post-processed according to wang et al. [Wang et al., 2022]. Each chemical can be represented as a graph G, which consists of nodes (atoms) and bonds (edges). The input to GCN layer is a set of node features, 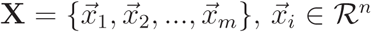, where n is the feature dimension of each node and m is the number of atoms. The propagation process can be calculated as follows:

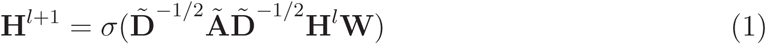

An adjacency matrix **Ã** ∈ *R*^*m×m*^ with self-connections represents the link relationships. 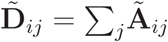 and **W** is the layer specific weight matrix. σ represents a activation function. **H**^*l*+1^ is the output of l + 1^*th*^ layer while **H**^0^ = **X**. DTSyn leveraged recitified linear unit (ReLU) function followed by GCN layer.

### 2.4 Gene embeddings

In the Protein-protein Interaction (PPI) network [Oughtred et al., 2019], nodes represent proteins (genes) and edges (PPI) indicate biological association among proteins. In order to obtain the numerical embeddings of proteins which contain PPI information, we used node2vec algorithm [Grover and Leskovec, 2016]. Since we used L1000 hallmark genes for expression profiles, we then selected L1000 gene representations for downstream analysis.

### 2.5 Cell line features

We extracted cell line expression profiles from CCLE [Ghandi et al., 2019]. The Library of Integrated Network-Based Cellular Signatures (LINCS) [Cheng and Li, 2016] proved that 978 hallmark genes can capture 80% information of the whole transcriptome. Therefore, we utilized hallmark genes as the initial cell line features. To further reduce the dimension of cell line features, we adopted a three-layer perceptron.

### 2.6 Coarse-granularity transformer for chemical-cell line and chemical-chemical associations

In DTSyn, two transformer-encoder blocks were adopted. DTSyn learned chemical-cell line and chemical-chemical associations via the coarse-granularity block. Multi-head attention is the core gradient of transformer [Wolf et al., 2020] which can model the interactions among chemicals and cell line. Specifically, the chemical feature vectors were obtained from the chemical feature matrix through pooling operation. We then concatenated chemical features with the densed cell line feature from MLP as an input. The coarse-granularity transformer block consists of two identical layers. Each layer has two sub-parts.The first is the multi-head attention mechanism and the second is feed-forward neural network **Figure 1**. The attention function is used to map a query and a set of key-value pairs to the output, where the query, key and value are all from input matrix. The attention scores can be calculated as follows:

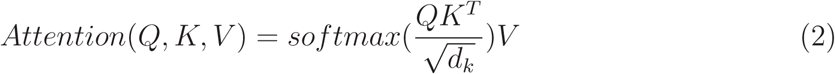

where Q, K, V are query, key and value respectively, d_*k*_ is the dimension of query and key. In DTSyn, each vector in the input (chemical, chemical, cell line) can be query, key and value.

In order to extract associations from different dimensions, we performed multi-head attention functions in parallel. On each of the attention module, we got different outputs. These outputs were concatenated resulting in the final values as follows:

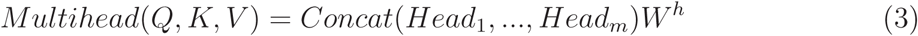

where 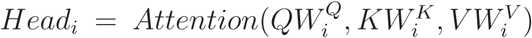, where *W*^*h*^, *W*^*Q*^, *W*^*K*^, *W*^*V*^ are learnable parameters.

In addition to multihead sub-layer, each transformer encoder contains a feed-forward neural network **Figure 1**. We applied two-layer linear transformation with ReLU as activation function. Coarse-transformer output can be generated as follows:

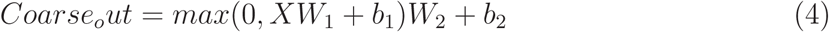

where *W*_1_, *b*_1_, *W*_2_, *b*_2_ are the parameters of linear functions.

### 2.7 Fine-granularity transformer for gene-chemical substructure, gene-gene associations

Identifying chemical-gene interactions (drug-target interactions) is a crucial step in drug discovery or drug repurposing [Hinnerichs and Hoehndorf, 2021]. Finding new targets of approved drugs also help to analyze and identify new drug combinations as well as desirable therapeutic effects. Gene-Gene interactions (Protein–protein interactions) are of pivotal importance in the regulation of biological systems and are consequently implicated in the development of disease states [Scott et al., 2016]. Thus, DTSyn utilized a transformer encoder to extract these associations. In this encoder, gene vectors and chemical atomic vectors were concatenated as the input. The gene vectors were obtained from node2vec algorithm, pre-trained on protein-protein interaction network [Jin et al., 2021]. The gene vectors and chemical atomic feature vectors were represented as query, key and values. The computing process of attention module and feed-forward module were the same as coarse-granularity transformer.

### 2.8 Predictions

After updating the numerical representations of chemical substructure and cell lines, an MLP was designed for prioritizing the synergistic drug pairs **Figure 1**. First of all, the outputs from two transformer blocks were flattened and concatenated as an input for MLP. The MLP consisted of two linear transformations with a ReLU activation in between.

## 3 Experimental setup

### 3.1 Data split strategies

We first conducted a 5-fold cross validation strategy. Four of folds were selected as training data and one fold was left as testing data. The hyper-parameters were selected through random split 5-fold cross validation. To test the performance under different situations, we further used different strategies. **Figure** 2 illustrated the other four cross validation strategies. To determine the generalization ability of our model, we left drugs and cells out of the training sets for novel drugs or novel cells predictions. In addition, we also splitted data based on drug pairs and tissue types.

**Figure 2:**
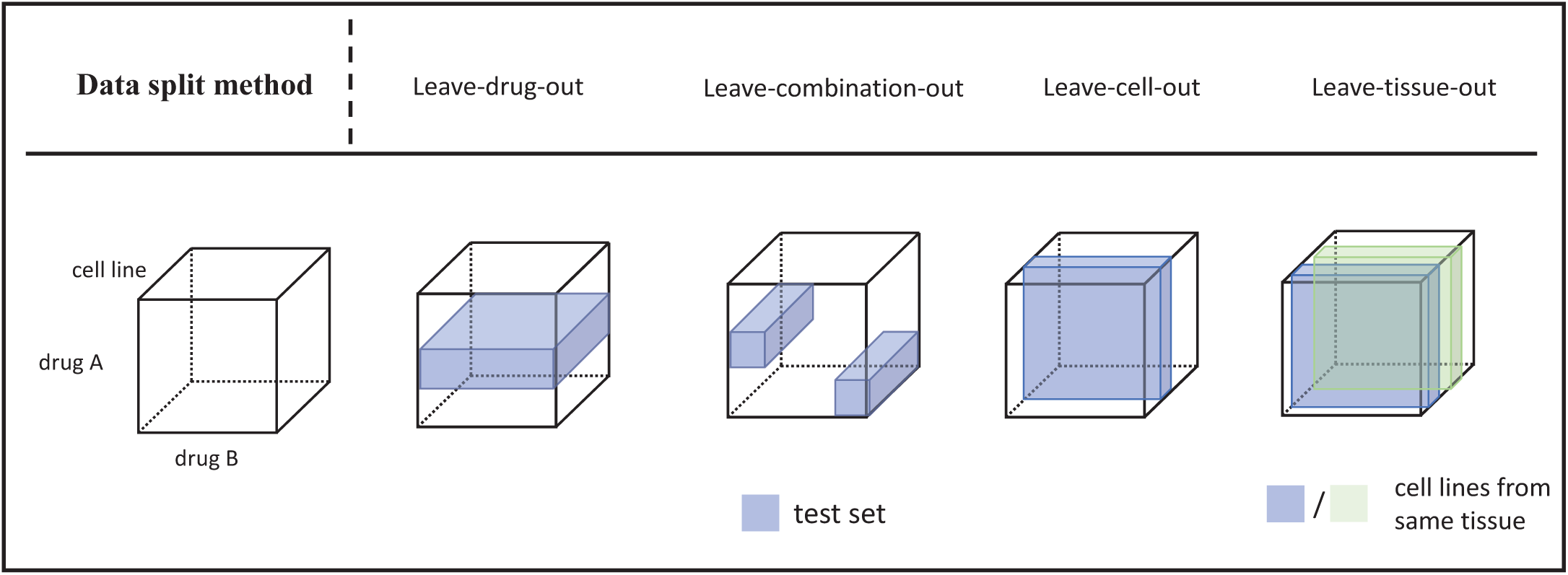
Four different data split strategies. The four splitting methods are shown in four columns. The blue color parts indicate testing data. The blue and green color parts in the fifth column represent different cell lines from the same tissue type.

### 3.2 Method comparisons

We compared DTSyn with other state-of-the-art (SOTA) deep learning methods and machine learning based methods on five splitting methods mentioned above. Three deep learning based methods were DeepDDs [Wang et al., 2022], DeepSynergy [Preuer et al., 2018] and three-layer MLP. The other machine learning based methods were random forest (RF) [Breiman, 2001], Adaboost [Freund and Schapire, 1997], SVM [Noble, 2006] and elastic net [Zou and Hastie, 2005]. The experiment results of both deep learning methods and machine learning methods were obtained from the same data input as DTSyn. Detailed settings for the compared methods were described in the **Supplementary Table S1**. To further compare the generalization ability of deep learning based methods, we employed five independent data sets mentioned above.

### 3.3 Global settings

In DTSyn, we set the input dimension of gene embedding as 128, the dimension of a cell line is 954 and the dimension of chemical atomic vector is 78. We used grid-search strategy to tune the optimal parameters of DTSyn. The hyperparameters of DTSyn were shown in **Table** 1. The hyper-parameters of DeepDDs and DeepSynergy were obtained from their original paper. Hyper-parameters of other competing methods were listed at **Supplementary Table S1**.

**Table 1:** Hyper-parameters of DTSyn

| Hyper-parameters | Values |
| --- | --- |
| Learning rate | 1e-2; 1e-3; 1e-4; 1e-5; **5e-6**; 1e-6 |
| GCN hidden size | [**512**, **128**]; [1024, 512, 128] |
| Pooling methods | **sum**; mean |
| Number of attention heads | 2; **4**; 8 |
| Dropout rate | 0.1; 0.2; 0.3; 0.4; 0.5; **0.6**; 0.7 |
| Activation function in transformer | **relu**; gelu |
| Hidden size in transformer | 32; **64**; 128 |
| Final FC hidden size | 2048; 1024; **512**; 256 |

The bold values represent the best performance.

### 3.4 Metrics

For the classification task of synergistic drug combinations, we adopted metrics including receiver operator characteristics curve (ROC-AUC),the area under the precision–recall curve (PR-AUC), accuracy (ACC), balanced accuracy (BACC), precision (PREC), True Positive Rate (TPR) and the Cohen’s Kappa value (KAPPA).

### 3.5 Independent data sets

We further applied DTSyn to predict five independent data sets that had not been tested previously. We also carried out experiments on novel drug pairs which do not exist in the training sets.

## 4 Results and analysis

### 4.1 Model comparisons

The comparison results of DTSyn and other competing methods on random split 5-fold cross validation were shown in **Supplementary Table S2**. DTSyn achieved ROC-AUC, PR-AUC, ACC, BACC, PREC, TPR and KAPPA of 0.89, 0.87, 0.81, 0.81, 0.84, 0.74 and 0.61, respectively. The result of random 5-fold cross validation of DTSyn was slightly inferior to results of DeepDDs. In addition, we validated the performance of DTSyn by switching the input order of two drugs. We compared the predicted labels given on two different input schemas. DTSyn achieved ROC-AUC, PR-AUC, ACC, BACC, PREC, TPR and KAPPA of 0.89, 0.88, 0.81, 0.81, 0.82, 0.78 and 0.61 after switching the input order. Obviously, drugs sequence did not affect the prediction capability of DTSyn.

The performance comparisons of four cross validation strategies were presented in **Table** 2. Notably, DTSyn achieved best performance on each cross validation task on ROC AUC. By leaving-drugs-out, DTSyn got a TPR of 0.65, which is 7% better than the second best methods (MLP) and 17% better than DeepDDs. As for leaving-combination-out task, DTSyn achieved TPR of 0.71, which is over 10% better than all competing methods. For leaving-cell-out task, the TPR of DTSyn is 0.75. DTSyn also achieved best results of PR AUC and TPR on leaving-tissue-out task. The performance comparison among DTSyn and other two deep learning methods on leaving-tissue-out task was shown in **Figure 3A**. DTSyn achieved best on all metrics except precision scores. Specially, DTSyn attained better results than DeepSynergy on ROC AUC and TPR with a strong evidence (*pvalue* ≤ 0.001). Further, DTSyn achieved better results than DeepDDs with moderate evidence (*pvalue* ≤ 0.05) on ACC, BACC and KAPPA, and with weak evidence (*pvalue* ≤ 0.1) on ROC AUC, PR-AUC, PREC and TPR. The performance on ROC AUC score of each deep learning method was illustrated in **Figure 3B**. Obviously, DTSyn achieved the best among all tissue types. For metric of TPR, DTSyn worked the best on colon, lung, breast, melanoma, ovarian and prostate (**Figure 3C**). Thus, DTSyn has potential to prioritize novel drug pairs across various tissue types.

**Table 2:** Performance comparisons on leave-drug-out, leave-combination-out, leave-cell-out and leave-tissue-out.

|  | Leave-drug-out |  |  | Leave-combination-out |  |  | Leave-cell-out |  |  | Leave-tissue-out |  |  |
| --- | --- | --- | --- | --- | --- | --- | --- | --- | --- | --- | --- | --- |
|  | ROC-AUC | PR-AUC | TPR | ROC-AUC | PR-AUC | TPR | ROC-AUC | PR-AUC | TPR | ROC-AUC | PR-AUC | TPR |
| DTSyn | **0.73 ± 0.02** | 0.70 ± 0.04 | **0.65 ± 0.19** | **0.78 ± 0.04** | 0.75 ± 0.06 | **0.71 ± 0.05** | **0.82 ± 0.02** | 0.79 ± 0.03 | **0.75 ± 0.04** | **0.81 ± 0.04** | **0.80 ± 0.03** | **0.74 ± 0.04** |
| DeepDDs | 0.71 ± 0.02 | 0.68 ± 0.04 | 0.48 ± 0.17 | 0.76 ± 0.02 | 0.75 ± 0.04 | 0.56 ± 0.09 | 0.81 ± 0.02 | 0.79 ± 0.03 | 0.69 ± 0.07 | 0.79 ± 0.04 | 0.79 ± 0.04 | 0.63 ± 0.10 |
| DeepSynergy | 0.66 ± 0.03 | **0.73 ± 0.03** | 0.46 ± 0.08 | 0.71 ± 0.02 | **0.77 ± 0.03** | 0.54 ± 0.06 | 0.75 ± 0.02 | **0.81 ± 0.03** | 0.60 ± 0.04 | 0.73 ± 0.04 | **0.80 ± 0.03** | 0.57 ± 0.09 |
| RF | 0.70 ± 0.02 | 0.67 ± 0.05 | 0.47 ± 0.19 | 0.73 ± 0.03 | 0.71 ± 0.05 | 0.57 ± 0.04 | 0.77 ± 0.03 | 0.75 ± 0.04 | 0.61 ± 0.06 | 0.79 ± 0.04 | 0.79 ± 0.04 | 0.63 ± 0.10 |
| Adaboost | 0.70 ± 0.02 | 0.67 ± 0.06 | 0.58 ± 0.13 | 0.71 ± 0.04 | 0.70 ± 0.04 | 0.62 ± 0.10 | 0.80 ± 0.01 | 0.78 ± 0.02 | 0.73 ± 0.03 | 0.80 ± 0.03 | 0.79 ± 0.02 | 0.68 ± 0.09 |
| SVM | 0.64 ± 0.07 | 0.62 ± 0.08 | 0.56 ± 0.11 | 0.68 ± 0.06 | 0.65 ± 0.07 | 0.62 ± 0.08 | 0.77 ± 0.03 | 0.75 ± 0.04 | 0.69 ± 0.03 | 0.76 ± 0.03 | 0.75 ± 0.03 | 0.66 ± 0.07 |
| MLP | 0.69 ± 0.08 | 0.68 ± 0.09 | 0.59 ± 0.21 | 0.72 ± 0.05 | 0.71 ± 0.07 | 0.63 ± 0.06 | 0.79 ± 0.03 | 0.77 ± 0.03 | 0.66 ± 0.05 | 0.76 ± 0.04 | 0.75 ± 0.04 | 0.62 ± 0.06 |
| Elastic net | 0.65 ± 0.06 | 0.63 ± 0.08 | 0.54 ± 0.21 | 0.68 ± 0.08 | 0.67 ± 0.08 | 0.60 ± 0.05 | 0.76 ± 0.02 | 0.74 ± 0.04 | 0.63 ± 0.03 | 0.75 ± 0.04 | 0.74 ± 0.03 | 0.62 ± 0.04 |

The bold values represent the best performance.

**Figure 3:**
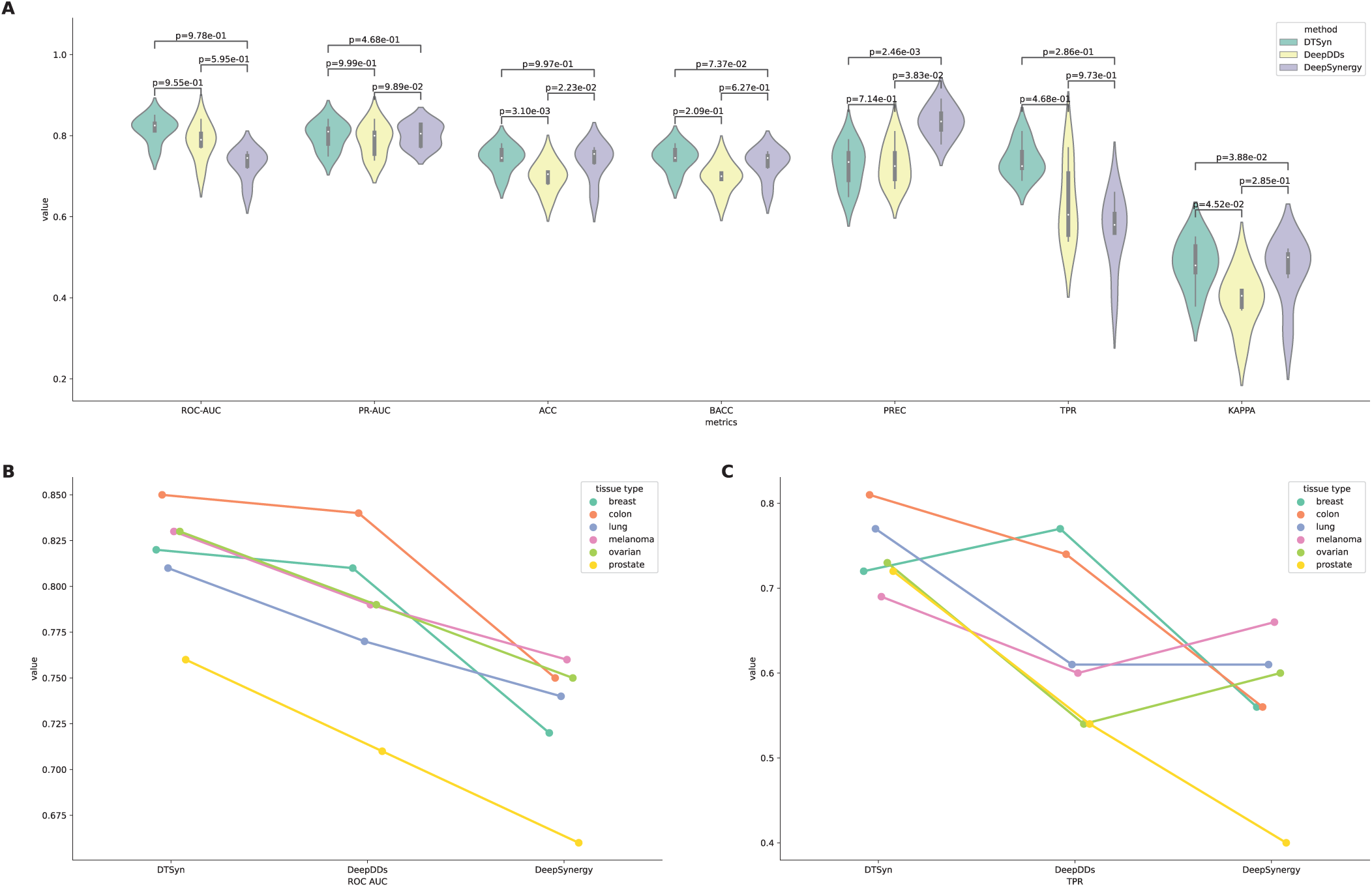
Model comparisons. **(A)**Comparison among three deep learning methods on seven metrics. **(B)**The ROC AUC value of three deep learning methods on each tissue type. **(C)**The PR AUC value of three deep learning methods on each tissue type.

### 4.2 Ablation study

To inspect the contribution of each transformer block in DTSyn, we thus designed three variants that consists of **DTSyn-C, DTSyn-F** and **DTSyn-B. DTSyn-C** is the model that only used fine-granularity transformer encoder. Without fine-granularity transformer block, the densed cell line feature was concatenated with output of fine-granularity transformer directly. **DTSyn-F** removed the fine-granularity transformer. The chemical atomic level features were concatenated with original gene embeddings without self-attention process. **DTSyn-B** is the variant that removes both transformer blocks. **Table** 3 summarized the results of the ablation study. Obviously, the performance of DTSyn-F was inferior to DTSyn, demonstrating that the chemical substructure-gene and gene-gene interactions multi-head attention can improve the performance of DTSyn. Further, by removing the coarse-granularity transformer block, DTSyn-C only achieved a ROC AUC of 0.71, indicating that the chemical-cell line transformer could extract the internal association for personalized medicines. Inaddition, DTSyn-F performed much better than DTSyn-C, presenting that the fine-granularity transformer block contributed more in our model. As for DTSyn-B, both transformer blocks were removed while only the feed-forward neural layers were used. DTSyn-B performed worst compared with other two variants. Based on the the performance comparison of these models, we concluded that two transformer blocks were of importance in our model and able to capture different aspects of interactions.

**Table 3:** Results of ablation study

| Methods | ROC-AUC | PR-AUC | ACC | BACC | PERC | TPR | KAPPA |
| --- | --- | --- | --- | --- | --- | --- | --- |
| DTSyn | **0.89 ± 0.01** | **0.87 ± 0.01** | **0.81 ± 0.01** | **0.81 ± 0.02** | **0.84 ± 0.02** | 0.74 ± 0.05 | **0.61 ± 0.03** |
| DTSyn-C | 0.71 ± 0.01 | 0.64 ± 0.01 | 0.67 ± 0.01 | 0.67 ± 0.01 | 0.64 ± 0.02 | 0.70 ± 0.04 | 0.34 ± 0.01 |
| DTSyn-F | 0.87 ± 0.01 | 0.84 ± 0.02 | 0.79 ± 0.01 | 0.79 ± 0.02 | 0.78 ± 0.03 | **0.77 ± 0.01** | 0.58 ± 0.02 |
| DTSyn-B | 0.69 ± 0.01 | 0.63 ± 0.01 | 0.66 ± 0.01 | 0.65 ± 0.01 | 0.66 ± 0.02 | 0.56 ± 0.02 | 0.30 ± 0.02 |

The bold values represent the best performance.

### 4.3 Experiments on independent datasets

Furthermore, we also evaluated the generalization performance of our model by testing on other five independent data sets. **Supplementary Figure S1** showed the predicted score distribution generated by DTSyn on five data sets. Since the data distribution was imbalance among these data sets, we paid more attention to BACC. As (**Figure**4A) shown, DTSyn achieved best on ALMANAC, FLOBAK, FORCINA and YOHE datasets with BACC of 0.57, 0.56, 0.53 and 0.48, respectively. Meanwhile, it achieved BACC of 0.51 on ASTRAZENECA dataset, which was slightly inferior than other two competing methods.

**Figure 4:**
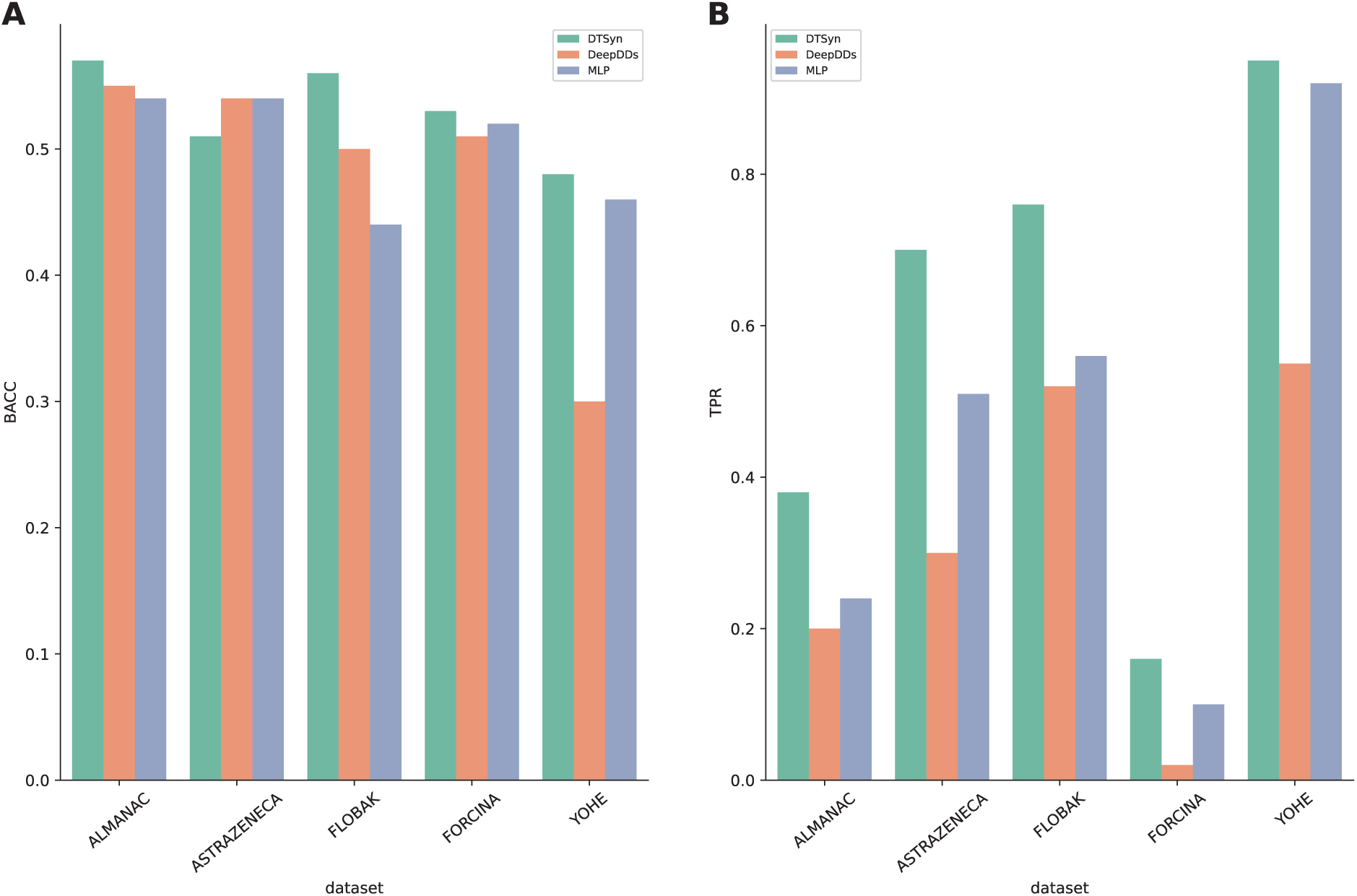
Indepedent datasets evaluation. **(A)**Model comparison based on BACC. **(B)**Model comparison based on TPR.

The detailed results were shown in **Supplementary Table S3**. We concluded that out model had better generalization ability, whereas those competing methods fell into overfitting.

### 4.4 Predictions on novel drug combinations

We further applied DTSyn to predict novel drug pairs that had not been tested previously. We combined all drugs and removed the existing drug pairs in the original training data, yielding 439 drug pairs in total. These novel drug combinations were tested on three typical cell lines (HCT116, HT29 and A375) [Lin et al., 2022]. **Figure**5) showed the distribution of all predicted probabilities on three cell lines. We compared the prediction performance of all cell lines and found prediction probabilities of A375 (melanoma) were significantly higher than those of two colorectal cancer (CRC) cell lines. We also evaluated the top predicted drug combinations across each cell line. **Supplementary Table S4** showed the top 10 predicted novel drug pairs on three cell lines.

**Figure 5:**
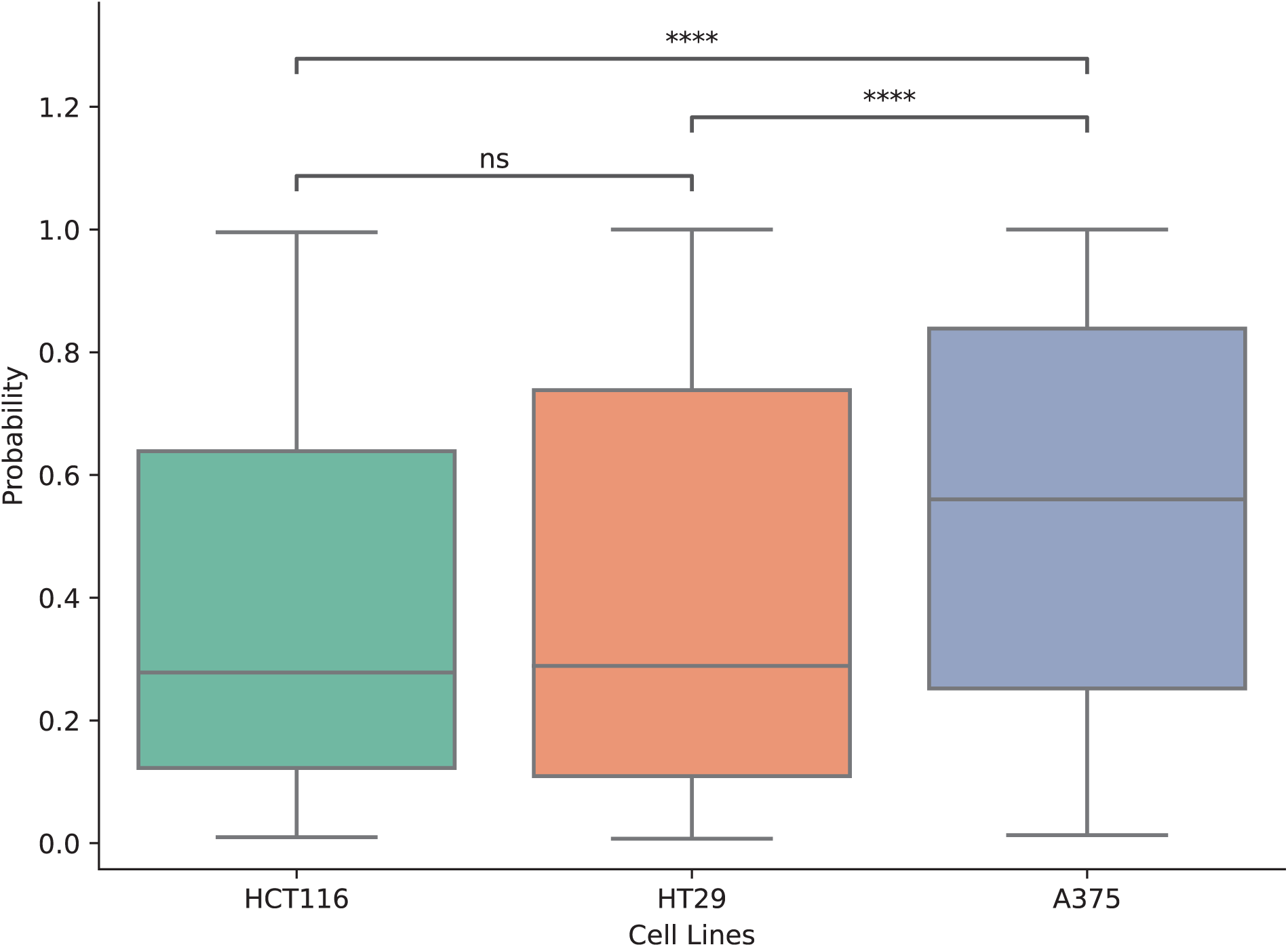
Predicted score on three cell lines. (ns: *pvalue* ≥ 0.1, ****: *pvalue* ≤ 1e − 4)

The combination of DINACICLIB and BEZ-235 achieved the predicted probability of 0.983 in HCT116 cell line. DINACICLIB is a potent, selective small molecule inhibitor of CDKs (CDK1, CDK2, CDK4, CDK5, CDK6 and CDK9) [Saqub et al., 2020]. It was reported that DINACICLIB acted against various human cancer cell lines [Parry et al., 2010]. In addition, CDK1 was proven to be a mediator of apoptosis resistance in BRAF V600E CRC [Li et al., 2019]. BEZ-235 ia a novel dual PI3K and mTOR inhibitor which has been widely tested in preclinical studies [Zhao et al., 2017]. Cretella et al. found that an orally-available inhibitor of CDK4/6 combined with PI3K/mTOR inhibitors impaired tumor cell metabolism in TNBC [Cretella et al., 2018]. Thus, DINACICLIB in combination with BEZ-235 may also be effective in HCT116 colon cell line.

We found that the combination of MK-8669 and METFORMIN had the highest prediction probability for A375 cell line. A375 is a human melanoma cell line and MK-8669 is a potent and selective mTOR inhibitor, preventing proliferation of a number of different tumor cell lines and xenografts [Rivera et al., 2011]. METFORMIN, which is a prescribed drug for type II diabetes, has been shown an great anticancer properties [Quinn et al., 2013]. It can activate adenosine monophosphate (AMP)-activated protein kinase (AMPK), which inhibits the mTOR signaling pathway [Mohammed et al., 2013]. A previous study suggested that the combination treatment with rapamycin (mTOR inhibitor) and METFORMIN synergistically inhibited the growth of pancreatic cancer (PC) in vitro and in vivo [Zhang et al., 2018].

HT29 is another colorectal cancer (CRC) cell line and MK-8669 and ZOLINZA may have the most potential in HT29. ZOLINZA, a hydroxamate histone deacetylase (HDAC) inhibitor, was particularly effective in inhibiting class I and II HDACs [Siegel et al., 2009]. Drug resistance emerges inevitably when mTOR inhibitor is used as single agent. The one of proposed escape pathways is the increasing phosphorylation of Akt, which is down regulated by HDAC inhibitors. Thus, the co-treatment with mTOR inhibitor and HDAC inhibitor may solve the resistance problem. It was concluded that patients with renal cell carcinoma experienced prolonged disease stabilization with HDAC and mTOR inhibitor combination treatment [Zibelman et al., 2015].

### 4.5 Explanation of transformer attention scores

As we expected, two transformer blocks can capture potential information of chemical substructure-gene interactions and chemical-cell line dependent associations. We analyzed the attention scores from the coarse-granularity transformer and fine-granularity transformer blocks. Here we used the drug combination of ETOPOSIDE and MK-2206 from CAOV3 and NCIH23 as an example. ETOPOSIDE was demonstrated to be an active chemotherapeutic drug used in neuroblastoma (NB) [Li et al., 2012]. MK-2206, an Akt inhibitor, binds to the Akt protein at pleckstrin-homology (PH) domain which leads to the conformation change of protein that prohibits its localization to plasma membrane thus deactivating its downstream pathways [Craig et al., 2007]. Investigation on the mechanisms underlying combination effect of ETOPOSIDE and MK-2206 showed that ETOPOSIDE-induced caspase-dependent apoptosis in neuroblastoma cells was enhanced when combined with MK-2206. Meanwhile, cell line-dependent mechanisms may also exist [Li et al., 2012]. The combination of ETOPOSIDE and MK-2206 showed a synergistic effect in CAOV3 (ovarian) cell line and an antagonistic effect in NCIH23 (lung) cell line. The significant attention scores between cell lines and two drugs may represent the effectiveness of each drug in that cell line. As shown in (**Figure**6), for CAOV3, the third column of each attention head obtained much higher attention scores compared to NCIH23, which means CAOV3 may benefit from the drug combination. In addition, each attention head may extract associations from different dimensions. To further investigated the interactions among drug pairs and potential interacted genes, we also analyzed the fine-granularity transformer attention scores. (**Figure**7) showed a part of the attention score heat map of combination of ETOPOSIDE and MK-2206 in CAOV3 of first attention head. The region with high association coefficients may indicate the chemical substructure-gene interactions. We observed that the genes SNAP25, GALE, PRKCD, PIK3R3 and DDIT4 had higher interaction coefficients. Since the atomic representations were obtained by two-layer GCN, thus each atom might represent chemical sub-structure. The previous study showed that synaptosomal-associated protein 25 (SNAP25), was associated with the effects of targeted chemotherapy [Huang et al., 2017]. Hodel et al. reported that the reduction of SNAP25 expression level providing a target for the development of therapeutic treatments [Hodel, 1998]. Further, SNAP25 mainly presented in the cytosol or recruited to the plasma membrane through the interaction with syntaxin (STX) proteins [Vogel et al., 2000]. A mechanism study illustarted that STX3 activated Akt-mTOR signaling to promote cancer proliferation, and Akt inhibitor MK-2206 repressed STX3 effects [Nan et al., 2018]. UDP-galactose-4-epimerase (GALE), a key enzyme of galactose metabolism,4,5 was overexpressed in some kinds of cancers, such as papillary thyroid carcinoma and glioblastoma [Sun et al., 2019]. Souza observed that GALE expression was associated with clinical-pathological parameters and the outcome of gastric adenocarcinoma patients [de Souza et al., 2019]. These evidence suggested that GALE may be a diagnostic biomarker and a potential therapeutic target. It was reported that inhibition of PRKCD protected kidneys during cisplatin treatment and enhanced the chemotherapy efficacy in tumors [Zhang et al., 2017]. PRKCD may suppress autophagy by phosphorylating AKT, which further phosphorylates MTOR to repress ULK1. Phosphoinositide-3-kinase regulatory subunit 3 (PIK3R3), a regulatory subunit of PI3K, which participated in tumor tumorigenesis and metastasis [Yoon et al., 2021]. The over expression of PIK3R3 in lung cancer was reported from wang et al.’s study [Wang et al., 2015]. Further, the inhibition of PIK3R3 can reverse the chemotherapy resistance [Wang et al., 2015]. We further investigated the interactions between drugs and DNA damage-inducible transcript 4 (DDIT4). The previous studies have illustrated that dysregulation of DDIT4 occured in various cancers with paradoxical roles [Fattahi et al., 2021]. Jin et al. reported that DDIT4 suppressed tumor through suppression of mTORC1 in non-small cell lung cancer [Jin et al., 2011]. While as an oncogene, upregulation of DDIT4 led to tumor proliferation, migration and invasion in-vivo [Schwarzer et al., 2005, Zeng et al., 2018]. It was reported that high expression level of DDIT4 was related to ovarian cancer (OC) [Chang et al., 2018]. Moreover, the expression level of DDIT4 can be upregulated by small molecules, such as dopaminergic neurotoxins and DNA damage agent ETOPOSIDE [Tirado-Hurtado et al., 2018]. Coronel et al. established p53-RFX7-DDIT4 as a signaling axis inhibiting mTORC2-dependent AKT activation [Coronel et al., 2022], which may be related to the effect of Akt inhibitor MK-2206. This example supported that DTSyn can provide reasonable clues for understanding the mechansims of drugs action. In addition, DTSyn has the potential to discover new biomarkers for different cancers.

**Figure 6:**
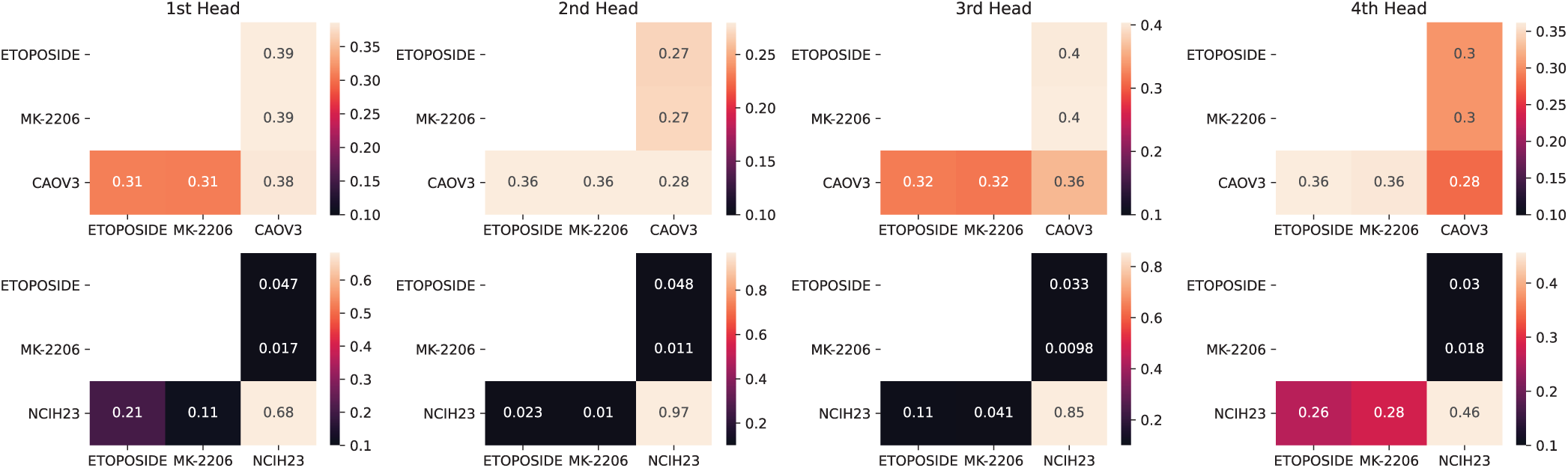
The heat maps of coarse-granularity transformer attention scores across ETOPOSIDE and MK-2206 on CAOV3 and NCIH23.

**Figure 7:**
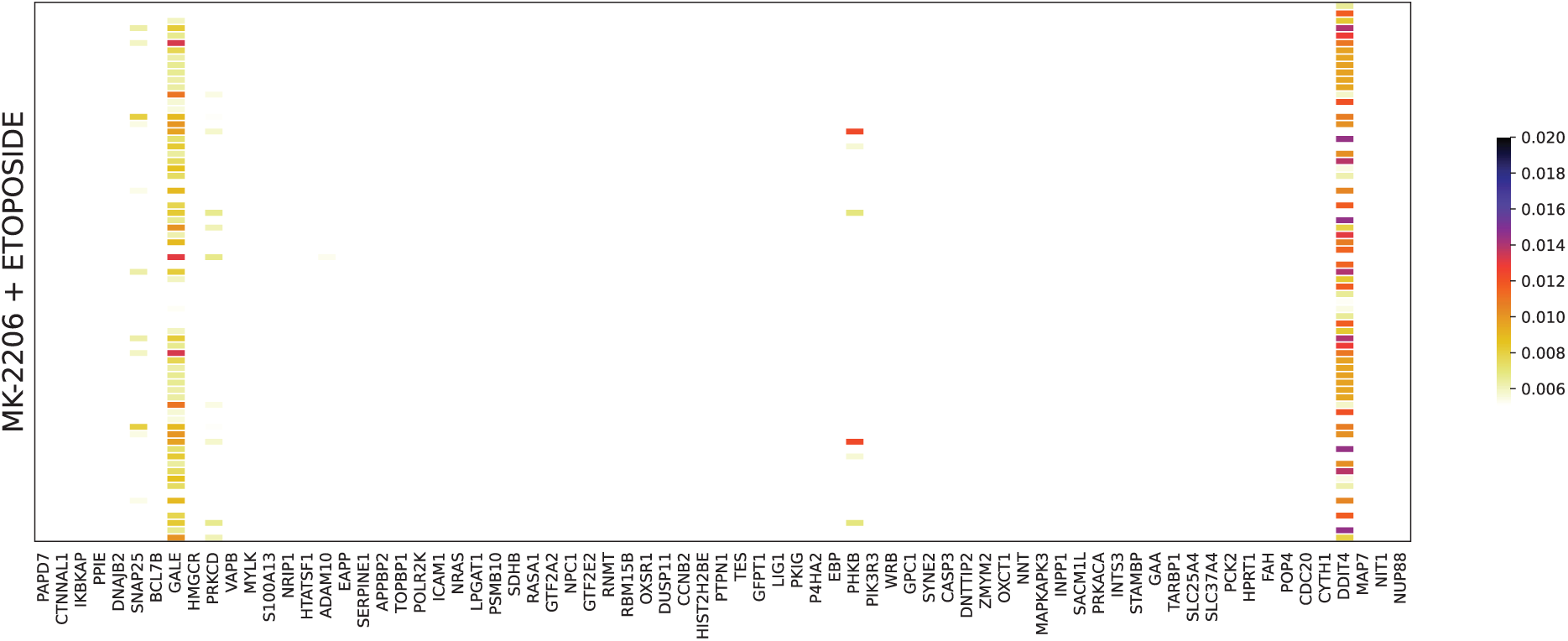
The heat map of fine-granularity transformer attention scores of ETOPOSIDE and MK-2206 on CAOV3.

We further analyzed the embeddings of chemicals after the coarse-granularity transformer and found that the synergistic and antagnostic drug combinations demonstrated obvious patterns. The comparison of drug combinations embeddings on three typical cell lines were shown in **Supplementary Figure S2**. A dimension reduction algorithm, UMAP [McInnes et al., 2018], was used to obtain the two-dimensional space of each drug pairs. After training, the synergistic combinations and antagnostic combinations fell into two obvious clusters. This further verified the effectiveness of DTSyn.

## 5 Conclusion and discussion

In this study, we proposed a novel dual-transformer based model, DTSyn, which can prioritize the potential drug combinations for cancer therapy. We utilized multi-head attention mechanism that can extract the chemical substructure-gene, chemical-chemical and chemical-cell line interactions. Specially, DTSyn is the first model which utilizes two transformer blocks to extract interactions relationships among genes, chemicals and cell line, which provides the understanding of mechanisms of drugs action. DTSyn models chemical-cell line associations through coarse-granularity transformer, which can extract relationships between expression level and chemical information. The embeddings of chemicals after this transformer can be clustered to two obvious groups. On the other hand, DTSyn uses another fine-granularity transformer to learn the associations among chemical-substructures and gene embeddings pretrained from PPI networks. This can help DTSyn to learn the potentail chemical-target interactions. Meanwhile, DTSyn has the ability to find cell line-dependent cancer-related genes which may play different roles in various cell lines under drug combinations. The ablation study also showed that two transformer blocks both contributed to the performance of DTSyn.

The performance of DTSyn was better than other comparative methods on four cross validation tasks while slightly inferior to DeepDDs on 5-fold cross validation. Besides, We were the first to conduct generalization experiments on five independent data sets never been seen before. Our model especially performed better than other two deep learning methods on TPR metric. Therefore, DTSyn has showed a better generalization ability when applied to different applications. Notably, switching the input order of drugs would not affect the performance of DTSyn.

Although DTSyn has demomstrated great performance, we noticed the balanced accuracy is still limited on independent data sets. The problem is expected to be solved by collecting of more training data. Another problem is that we only used 978 hallmark genes to train the fine-granularity transformer, which may lose some chemical-target interactions. Besides, DTSyn only used expression data as features for cell line. However, other omics data, such as methylation and genetic data can be included in the future, which may help in representing the cell line systematically.

In conclusion, our study suggests that DTSyn utilizing dual-transformers has the great potential in identifying novel synergistic drug pairs and also providing possible interpretability in mechanisms of drug actions.

## 7 Author contributions statement

J.H. conceived the experiment(s), J.H. and X.F. conducted the experiment(s), J.H. and J.G. analysed the results. J.H., F.W., Z.L. and G.Z. wrote and reviewed the manuscript.

