## Supplemental tables and figures for "DTSyn: a dual-transformer-based neural network to predict synergistic drug combinations"

### S1 Methods

#### Hyper-parameters in Global setting

The hyper-parameters of all competing methods were obtained based on five-fold cross validation method. The competing methods are Random Forest, Adaboost, SVM, MLP and Elastic net. Table S1 lists all hyper-parameters of each comparative algorithms.

| Methods | hyper-parameters | values considered |
|---|---|---|
| Random Forest | number of estimators | 128; 512; **1024**; 2048 |
| Adaboost | number of estimators<br>learning rate | 100; **200**; 300; 400; 500; 600<br>**1**; 1e-1; 1e-2; 1e-3; 1e-4 |
| SVM | C<br>tolerance for stopping criterion | 1e-4; 1e-3; 1e-2; 1e-1; **1**; 10; 100<br>0.1; **0.05**; 0.025; 0.005; 0.001 |
| MLP | number of hidden layer<br>pooling methods<br>hidden size | 1<br>mean; **sum**<br>512; 1024; 2048; **4096** |
| Elastic net | C<br>l1 ratio | 0.01; **0.1**; 1; 10; 100<br>0.25; **0.5**; 0.75 |

Table S1: Hyper-parameters and corresponding values for competing methods

### S2 Results

#### Performance comparison on 5-fold cross validation

To obtain the best parameters of DTSyn and other comparative methods, we performed 5-fold cross validation to choose the optimal parameters. The prediction distribution of DTSyn is presented in Figure S1. The detailed comparative results are represented in Table S2.

| Methods | ROC-AUC | PR-AUC | ACC | BACC | PREC | TPR | KAPPA |
|---|---|---|---|---|---|---|---|
| DTSyn | $0.89 \pm 0.01$ | $0.87 \pm 0.01$ | $0.81 \pm 0.01$ | $0.81 \pm 0.02$ | $\mathbf{0.84 \pm 0.02}$ | $0.74 \pm 0.05$ | $0.61 \pm 0.03$ |
| DeepDDs | $\mathbf{0.91 \pm 0.01}$ | $\mathbf{0.90 \pm 0.01}$ | $\mathbf{0.83 \pm 0.01}$ | $\mathbf{0.83 \pm 0.01}$ | $0.82 \pm 0.02$ | $\mathbf{0.82 \pm 0.03}$ | $\mathbf{0.66 \pm 0.02}$ |
| DeepSynergy | $0.72 \pm 0.01$ | $0.77 \pm 0.03$ | $0.72 \pm 0.01$ | $0.72 \pm 0.01$ | $0.73 \pm 0.05$ | $0.64 \pm 0.02$ | $0.43 \pm 0.02$ |
| RF | $0.74 \pm 0.03$ | $0.73 \pm 0.03$ | $0.67 \pm 0.01$ | $0.67 \pm 0.02$ | $0.70 \pm 0.07$ | $0.59 \pm 0.03$ | $0.35 \pm 0.04$ |
| Adaboost | $0.74 \pm 0.02$ | $0.72 \pm 0.03$ | $0.75 \pm 0.02$ | $0.66 \pm 0.02$ | $0.63 \pm 0.08$ | $0.69 \pm 0.08$ | $0.32 \pm 0.04$ |
| SVM | $0.68 \pm 0.05$ | $0.65 \pm 0.06$ | $0.62 \pm 0.05$ | $0.62 \pm 0.05$ | $0.59 \pm 0.05$ | $0.66 \pm 0.06$ | $0.25 \pm 0.09$ |
| MLP | $0.84 \pm 0.01$ | $0.82 \pm 0.01$ | $0.76 \pm 0.01$ | $0.75 \pm 0.01$ | $0.75 \pm 0.01$ | $0.71 \pm 0.01$ | $0.50 \pm 0.02$ |
| Elastic net | $0.68 \pm 0.08$ | $0.67 \pm 0.07$ | $0.63 \pm 0.07$ | $0.63 \pm 0.07$ | $0.61 \pm 0.08$ | $0.62 \pm 0.07$ | $0.27 \pm 0.14$ |

The bold values represent the best performance.

Table S2: Comparison on 5-fold cross validation

#### Results of independent datasets

In order to validate the generalization ability of DTSyn, we applied DTSyn to five different datasets. The detailed results of DTSyn and other two competing deep learning methods are listed as Table S3.

#### Prediction results on novel drug pairs

The novel drug pairs which were not combined in the training data were tested on three typical cell lines (HCT116, HT29 and A375). We listed top 10 predicted drug combinations based on probability scores in Table S4.

| Methods | dataset | ACC | BACC | TPR |
| --- | --- | --- | --- | --- |
| DTSyn | ALMANAC | 0.63 | 0.57 | 0.38 |
| | ASTRAZENECA | 0.56 | 0.51 | 0.7 |
| | FLOBAK | 0.49 | 0.56 | 0.76 |
| | FORCINA | 0.29 | 0.53 | 0.89 |
| | YOHE | 0.89 | 0.48 | 0.95 |
| DeepDDs | ALMANAC | 0.63 | 0.55 | 0.20 |
| | ASTRAZENECA | 0.51 | 0.54 | 0.30 |
| | FLOBAK | 0.45 | 0.50 | 0.52 |
| | FORCINA | 0.22 | 0.51 | 0.02 |
| | YOHE | 0.56 | 0.30 | 0.55 |
| MLP | ALMANAC | 0.62 | 0.54 | 0.24 |
| | ASTRAZENECA | 0.57 | 0.54 | 0.51 |
| | FLOBAK | 0.38 | 0.44 | 0.56 |
| | FORCINA | 0.0.30 | 0.52 | 0.10 |
| | YOHE | 0.56 | 0.46 | 0.92 |

Table S3: Comparison on five independent datasets

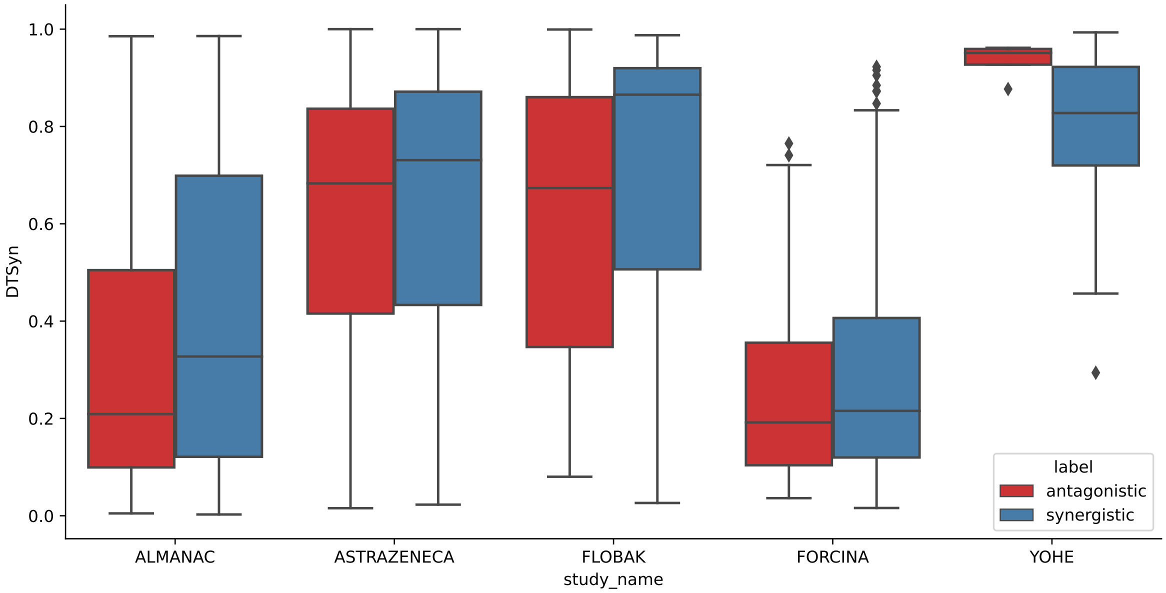


Figure S1: Prediction distribution of DTSyn on five independent datasets.

#### Analysis of embeddings of two chemicals

In order to explore the effectiveness of coarse-granularity transformer, we visulized the embeddings of chemicals before and after the coarse-granularity transformer. We also used three typical cell lines (HCT116, HT29 and A375) as examples. We adopted UMAP algorithm to obtain two-dimentional space of chemicals. After training, the synergistic pairs and antagnostic pairs fell in to two obvious clusters.

| Chemical1 | Chemical2 | Cell line | probability |
|---|---|---|---|
| MK-8669 | ZOLINZA | HT29 | 1.0 |
| MK-8669 | METFORMIN | A375 | 0.999 |
| MK-5108 | DEXAMETHASONE | A375 | 0.999 |
| CARBOPLATIN | MK-4541 | HCT116 | 0.996 |
| OXALIPLATIN | BEZ-235 | A375 | 0.987 |
| MK-2206 | ETOPOSIDE | A375 | 0.986 |
| DINACICLIB | BEZ-235 | HCT116 | 0.983 |
| BEZ-235 | DASATINIB | HT29 | 0.978 |
| LAPATINIB | PACLITAXEL | HT29 | 0.977 |
| MK-8669 | VINBLASTINE | HT29 | 0.976 |

Table S4: Top 10 predicted drug combinations on three typical cell lines.

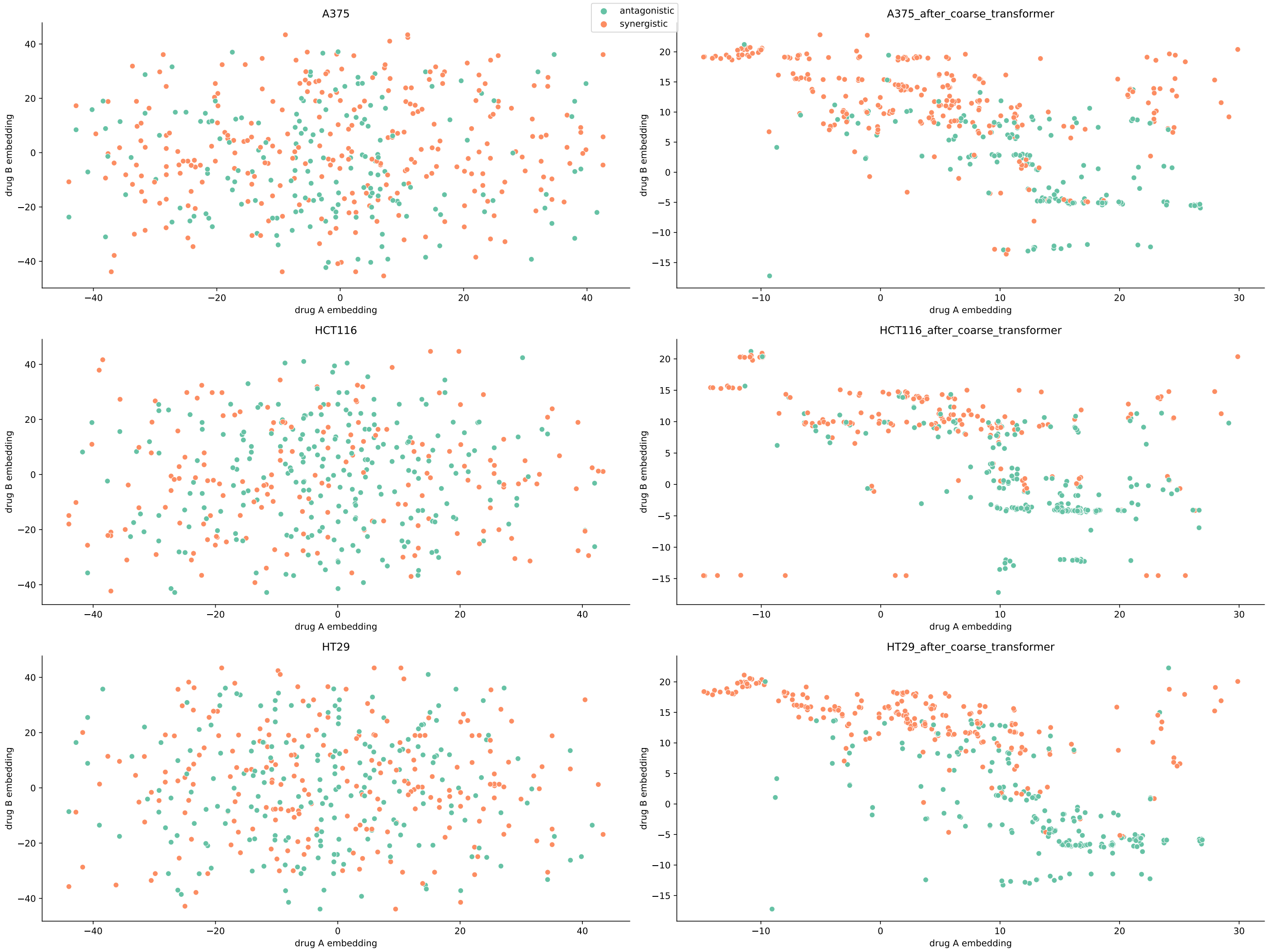


Figure S2: Scatter plots of chemical embeddings before and after training on three typical cell lines.
